# Palliative care for people living with HIV/AIDS: Factors influencing healthcare workers’ knowledge, attitude and practice in public health facilities, Abuja, Nigeria

**DOI:** 10.1101/460709

**Authors:** Whenayon Simeon Ajisegiri, Aisha A. Abubakar, Abdulrazaq A. Gobir, Muhammad S. Balogun, Kabiru Sabitu

## Abstract

Physicians, nurses and allied health staff play very vital roles in addressing palliative care (PC) needs of people living with HIV/AIDS (PLWHA). The healthcare professionals’ experiences determine the success of palliative care delivery. There is paucity of data on palliative care for PLWHA. For this reason, we assessed the knowledge, attitude and practice of palliative care for PLWHA and associated factors among health care professionals.

We conducted a cross-sectional descriptive study among healthcare workers (HCWs) in public health facilities in the Federal Capital Territory, Nigeria between February and May, 2017. Multistage sampling technique with proportionate-to-size allocation was used to determine facility sample size and HCWs per professional discipline. Data were collected with questionnaires adapted from Palliative Care Quiz for Nursing, Frommelt Attitude toward Care of the Dying and practical questions adapted from PC standard guidelines. Univariate analysis was done to compute frequencies and proportions. Odd’s ratios were calculated to assess the statistical association between variables and P-value ≤ 0.05 was considered significant. Multivariate analysis was conducted on variables that were significant with bivariate analysis. Epi-Info software was used for statistical analyses.

The mean age of the 348 participants was 37.5 years (SD: ±8.9) and 201 (57.8%) were female. Thirty-six (10.3%) participants had good knowledge, 344 (98.9%) had favourable attitude and 90 (25.9%) had good practice of PC for PLWHA. Major determinants of good knowledge of PC include being a doctor (aOR = 2.70; 95%CI: 1.28 - 5.56). Determinants of good practice of palliative care include possessing a minimum of a bachelor degree (aOR=2.30; 95%CI : 1.05-5.08) and practicing in a tertiary hospital (aOR=6.67; 95%CI : 3.57-12.5).

HCWs had favourable attitude towards PC for PLWHA despite an overall poor knowledge and practice. We recommended quality in-service training and continuous education on palliative care for HCWs.

## Introduction

Sub-Saharan Africa is the most affected region in terms of Human Immunodeficiency Virus (HIV) infection ad it harbours about 70% of the global population of people living with HIV/AIDS (PLWHA)[1, 2]. Nigeria, being the most populous country in Africa with estimated population of 182 million[3] has an estimated national HIV prevalence rate of 3.4%.[4] and that of the Federal Capital Territory, Abuja is 7.5%[5].

The advent of highly active anti-retroviral therapy (HAART) has transformed HIV infection and AIDS to another chronic illness that can be managed but not cured. Despite the use of HAART, PLWHA still continue to die at a higher rate than the uninfected individuals[6]. These among other issues necessitated the need for palliative care as part management of HIV/AIDS[6, 7].

“Palliative care is an approach which improves the quality of life of patients and their families facing life-threatening illness, through the prevention, assessment and treatment of pain and other physical, psychosocial and spiritual problems”[8]. It provides pain relief, integrates psychological and spiritual aspects of patients care and offers support system to both the patients and their relatives to live an active live and cope during patient’s illness/death respectively[9]. Very few researches have been conducted to explore the activities and impact of palliative care for HIV as most attention is placed on ART use[10].

Physicians, nurses and allied health staff play very vital roles in addressing the physical, social and spiritual needs of PLWHA. The health care professionals’ knowledge, attitude, belief and experience go a long way to determine how successful the delivery of palliative care will be[11,12]. In order to strengthen the continuum of care for PLWHA, there is need for considerable investment into research and education on palliative care for health workers who provide services to these population[13]. There is paucity of data on healthcare workers’ knowledge and practice with regards to palliative care for PLWHA. For this reason, we assessed the knowledge, attitude and practice of health care professionals towards palliative care for PLWHA.

## Methods

**Study design, area & period**: We conducted a cross-sectional descriptive study among healthcare workers (doctors and nurses) in public secondary and tertiary health facilities in Abuja, in the Federal Capital Territory (FCT), between February and May, 2017. Abuja is the capital city of Nigeria with a population of 1,406,239. The FCT consists of six area councils and a total of 293 hospitals offering HIV/AIDS services. Of these, government-owned facilities comprised 189 primary, 14 secondary and 3 tertiary health facilities. The health workers all consist of 1,137 doctors, 2,549 nurses and about 900 other allied health staff that offer testing, treatment and prevention services to PLWHA.

We determined the sample size using a prevalence (p) of 30.5% [13], a degree of precision of 5%, a standard normal deviate (Za) of 1.96. Using the formula for descriptive health studies n=,Z^2^_a_ pq/d^2^[14]; this gave us 326. The estimated minimum sample size was 333 after accounting for 10% non-response rate and correcting for finite population. Total population of doctors and nurses in FCT Health sector is 3686.

**Sampling technique:** A multistage sampling techniques was used. Proportionate allocation sampling based on each of the 17 health facilities’ staff strength was used to determine facility sample size. A proportionate-to-size sampling was further used to determine the number of HCWs selected per professional discipline

**Study population:** Eligible participants were doctors and nurses directly involved in the management of PLWHA or provision of any HIV/AIDS-related services in the public secondary and tertiary health facilities. Allied health staff were excluded.

Five research assistants fluent in speaking English language and with previous experience in health-related research activities were recruited and trained for a day to ensure standards. All respondents understand and speak English language and so, there was no need for to translate to local language. Data was collected for a period of three months.

**Data collection method, tool and operational definitions:** Data were collected using a self-administered questionnaire on “Open Data Kit (ODK Collect)” application on Android mobile phones. The data collection instrument contains four sections. **Section A:** Health workers’ socio-demographic characteristic: age, gender, hospital, qualification, job position, department of work, working experience, training in palliative care. **Section B:** Participants' knowledge was assessed using questions adapted from Palliative Care Quiz for Nursing (PCQN) [13]. One (1) point was awarded for each correct answer while “incorrect” or “not sure” answers took a zero (0) score. Correct responses were summed up to get a total knowledge score for each participant. The knowledge score was classified into Poor knowledge (<75%) and Good knowledge (≥75%)[13]. **Section C:** Attitude was assessed using a 5-item Likert scale with questions adapted from Frommelt Attitude toward Care of the Dying (FATCOD) [13]. It represented participant’s attitudes to a subject scored on 5-point scale - 1 (Strongly Disagree), 2 (Disagree), 3 (Uncertain), 4 (Agree) to 5 (Strongly Agree). Some questions were worded positively and others negatively. The score of negative items were reversed to calculate the attitude. Overall score was calculated by adding each individual’s scores out of 90. The attitude score was categorized into favourable (≥50%), and unfavourable (<50%). A higher score represented a more positive attitude toward palliative care[13]. **Section D:** Practice was assessed through 11 practical questions adapted from previous studies, standard guidelines and literatures related to PC practice. The practice scores were categorized into good practice (≥75%) and poor practice (<75%)[13].

**Data analysis:** Data was exported from ODK as Microsoft Excel 2007 version, cleaned and entered into Epi Info version 7 for statistical analyses. Univariate analyses: descriptive statistics was done to describe frequencies and proportions. Bivariate analysis was done and odd’s ratios were computed to assess the statistical association between independent and dependent variables. Variables were considered significant at a P-value ≤ 0.05. Multivariate analysis was conducted on variables that were significant with bivariate analysis.

**Ethical approval:** Ethical approval was obtained from the Ethics and Research Committee of the Federal Capital Authority Administration and the selected Health Facilities. Verbal consent was obtained from each respondent before administering questionnaire. Participants’ confidentiality and anonymity was maintained all through the process of the research. They were also informed that they could withdraw from the study at any stage they desire without any consequence.

## Results

The mean age of the 348 participants was 37.5 years (Standard Deviation: ±8.9), and about 43.7% of the participants were within age-group 30-39 years. Majority, 201 (57.8) were female and 274 (78.7) were married, 222 (63.8) were nurses, 231 (66.4%) possess a minimum of bachelor degree and 230 (66.0%) have work experience of 10 years or less. About 181 (52.5%) of respondents said their undergraduate curriculum consists of palliative care while 70 (20.3%) where not sure. (Table 1)

**Table 1.** Socio-demographic profile of healthcare workers at public secondary and tertiary hospitals Abuja, May 2017

| Socio-demographic variables | Frequency<br>(n = 348) | Percentage (%) |
| --- | --- | --- |
| **Age (years)** |  |  |
| $\leq 29$ | 64 | 18.4 |
| 30-39 | 152 | 43.7 |
| 40-49 | 88 | 25.3 |
| 50-59 | 44 | 12.6 |
| **Mean age (SD): 37.5 years ($\pm 8.9$)** |  |  |
| **Sex** |  |  |
| Male | 147 | 42.2 |
| Female | 201 | 57.8 |
| **Profession** |  |  |
| Doctor | 126 | 36.2 |
| Nurse | 222 | 63.8 |
| **Marital Status** |  |  |
| Married | 274 | 78.7 |
| Unmarried | 74 | 21.3 |
| **Highest of academic qualification** |  |  |
| Diploma (ordinary & higher) | 117 | 33.6 |
| Bachelor Degree | 138 | 45.4 |
| Post-graduate | 73 | 21.0 |
| ***Length of post-qualification work experiences (years)*** |  |  |
| ≤ 10 years | 230 | 66.9 |
| > 10 years | 118 | 33.1 |
| ***Undergraduate curriculum consists of palliative care (n =308)*** |  |  |
| Yes | 181 | 52.5 |
| Not sure | 70 | 20.3 |
| No | 94 | 27.2 |
| ***Received formal in-service training on palliative care (n=348)*** |  |  |
| Yes | 112 | 36.7 |
| No | 246 | 63.3 |

Only few 36 (10.3%) participants had good knowledge of palliative care. The minimum and maximum scores out of 20 were 1 and 19 respectively with an average score of 11.1 (SD: 3.21). Majority of the participants, 310 (89.1%) agrees that palliative care focuses on the relief and prevention of suffering and 319 (91.7%) believe that PLWHA requires palliative care. However, majority of the participants have some misconceptions about palliative care. First of this misconception is that, 252 (72.6%) believes “palliative care is disease-oriented and not person oriented”. Others include the belief that palliative care “is concerned with prolongation of life”, 279 (80.6%); “use of placebos is appropriate in the treatment of some types of pain”, 252 (72.6%); extent of the disease determines the method of pain treatment, 283 (81.6%) and that palliative care is just terminal care, 165 (47.4%). Majority agreed that “palliative care incorporates the whole spectrum of care: medical, nursing, psychological, social, cultural and spiritual” 285 (81.9%) and that “patient’s family members’ views should be considered in palliative care, 297 (85.6%)” (Table 2).

**Table 2.** Health Care Workers’ knowledge towards palliative care in public secondary and tertiary hospitals, Abuja, May 2017 (n=348)

| **Knowledge area assessed on palliative care** | **Right option,**<br>**N (%)** |
| --- | --- |
| Focus of palliative care | 310 (89.1) |
| Palliative care orientation | 70 (20.2) |
| Palliative care on prolongation of life. | 55 (15.9) |
| Use of multidisciplinary or inter-professional approach | 312 (89.7) |
| Incorporation of whole spectrum of care | 285 (81.9) |
| Identifying of the underlying causes for pain management | 284 (81.8) |
| Primary concern of palliative care | 122 (35.1) |
| Differentiation of palliative Care from hospice Care | 134 (38.8) |
| Differentiation of palliative Care from terminal Care | 143 (41.1) |
| Timing of palliative care institution | 140 (40.5) |
| Method of pain treatment | 32 (9.2) |
| Use of adjuvant therapies palliative care pain management | 277 (79.8) |
| Use of placebos for pain management | 252 (72.6) |
| Involvement of family members in palliative care | 297 (85.6) |
| Palliative care for PLWHA | 319 (91.7) |
| Focus of palliative care | 310 (89.1) |
| Palliative care orientation | 70 (20.2) |
| Palliative care on prolongation of life. | 55 (15.9) |

Of the 348 participants, 344 (98.9%) had favourable attitude towards palliative care for PLWHA. The minimum and maximum scores out of 90 were 38 and 81 respectively with an average score of 61.1 (SD: 6.9). About 52% of the participants disagreed (87{25.1%} and 83 {26.8%} strongly disagreed) that palliative care should be given only for dying PLWHA while 317 (91.1%) agreed that family should be involved in the physical care of the dying PLWHA. One hundred and fifty-nine (45.7%) participants agreed that they would be uncomfortable talking about death with a dying PLWHA. (Table 3).

**Table 3.** Attitude of healthcare workers towards palliative in public secondary and tertiary hospitals, Abuja, May 2017 (n=348)

| Statements on attitude of health care workers on palliative care | Agreement, n (%) |
| --- | --- |
| Palliative care should be given only for dying PLWHA | 181 (52.2) |
| As a patient nears death; the HCW should withdraw from his/her involvement with the patient | 277 (80.3) |
| The length of time required to give care to dying PLWHA would frustrate me | 182 (54.3) |
| Family should maintain as normal an environment as possible for their dying member. | 15 (4.3) |
| The family should be involved in the physical care of the dying PLWHA. | 18 (5.2) |
| It is not difficult to form a close relationship with the family of a dying PLWHA. | 62 (17.8) |
| HCW's care for the patient's family should continue throughout the period of grief and bereavement | 53 (15.2) |
| HCW's care should be extended to the family of the dying PLWHA. | 46 (13.2) |
| I am afraid to become friends with PLWHA | 238 (68.8) |
| I would be uncomfortable if I entered the room of a terminally ill HIV/AIDS person and found him/her crying | 121 (34.8) |
| Giving professional care to the chronically sick patient is a worthwhile learning experience | 21 (6.0) |
| Family members who stay close to a dying person often interfere with a professionals' job with the patient. | 67 (19.4) |
| HCW should not be the one to talk about death with the dying person | 100 (29.8) |
| The dying person and his/her family should be the in-charge decision makers | 79 (23.0) |
| I would be uncomfortable talking about impending death with the dying Person | 112 (32.2) |

Good practice of palliative care for PLWHA among the healthcare workers was 25.9%. There is a weak positive correlation between knowledge and practice of palliative care which was found to be statistically significant [r = +0.26, p = 0.001]. Majority of the participants, 292 (84.1%) initiates palliative care discussion during patients’ diagnosis while 290 (83.6%) informs terminally-ill patients about their diagnosis. Medical 263 (84.6%) and cultural 94 (30.2%) factors were respectively the most and least considered factors by participants when dealing with terminally ill patients, conditions. Regarding psychological issues, 22 (6.3%) participants hide the truth from the patients, 301 (86.5%) prefer to counsel them while 196 (56.3%) provide emotional support to the patients. Morphine 240 (69.0%) and Pentazocine 194 (55.7%) were the most commonly used drugs for treatment of severe pain by participants across all centres. Majority, 309 (88.8%) perceived the concerns raised by PLWHA or terminally ill patient as their right to treatment while 39 (11.2%), perceived it as threat (Table 4).

**Table 4.** Practice of healthcare workers towards palliative care at public secondary and tertiary hospitals, Abuja, Nigeria-May, 2017

| Practice of palliative care | Options | Agreement<br>N (%) |
| --- | --- | --- |
| Initiate palliative care discussion | During diagnosis | 292 (84.1) |
|  | When the disease progress | 146 (42.1) |
|  | At the end of life | 18 (5.2) |
| Inform terminally-ill patient/PLWHA about their diagnosis | Yes | 290 (83.6) |
|  | No | 13 (3.7) |
|  | Depending on family's wish | 175 (50.4) |
| Factors considered when dealing with terminally-ill patient/PLWHA | Spiritual | 140 (45.0) |
|  | Medical condition | 263 (84.6) |
|  | Cultural | 94 (30.2) |
|  | Psychological | 148 (47.6) |
| Ways by which spiritual issues are addressed | Connect with spiritual counsellor | 225 (65.0) |
|  | Impose your own view | 53(15.3) |
|  | Understand patient reaction | 107 (30.9) |
| Ways by which psychological issues addressed | Emotional support | 196 (56.3) |
|  | Counselling the patient | 301 (86.5) |
|  | Hiding the truth | 22 (6.3) |
| Cultural issues to be considered during patient care | Truth telling and decision making | 194 (56.4) |
|  | Language | 128 (37.2) |
|  | Family communication | 213 (61.9) |
|  | Perspective on grieving, suffering & death | 102 (29.3) |
| People involved during decision making | Patient | 269 (77.3) |
|  | Family | 279 (80.2) |
|  | Other health professionals | 196 (56.3) |
|  | My decision alone | 12 (3.4) |
| Perception of concerns or questions raised by PLWHA or terminally ill patient | Patient's right | 309 (88.8) |
|  | Threat | 39 (11.2) |
|  | Attention-seeking behaviour | 103 (29.6) |
|  | Doubting your professionalism | 13 (3.4) |
| Communication with the family of PLWHA/terminally ill patient depends on: | Family's ability to assimilate information | 266 (76.9) |
|  | Their involvement in decision making | 243 (70.2) |
|  | Your willingness to disclose information | 46 (13.3) |
| Commonly use medication for severe pain | Paracetamol | 94 (27.0) |
|  | Ibuprofen | 112 (32.2) |
|  | Morphine | 240 (69.0) |
|  | Codeine | 95 (27.3) |
|  | Pentazocine | 194 (55.7) |

Doctors were more knowledgeable on Palliative care for HIV than nurses (OR = 2.1414; 95% CI: 1.07 - 4.29; P = 0.03). HCWs who worked in secondary health facilities were also found to be more knowledgeable than those working in tertiary health facilities (OR= 3.67; 95% CI: 1.63 - 8.33; P = 0.01). On the other hands, those who had in-service training (OR = 0.47; 95% 0.23 - 0.97; P = 0.0379) and those whose undergraduate curriculum contain palliative care module (OR = 0.27; 95% CI: 0.10- 0.72; P = 0.01) were respectively found to be less knowledgeable when compared with those who did not. With regards to factors associated with attitude towards palliative care, those who were married had a more favourable attitude towards Palliative care for HIV patients the unmarried (single, widowed or separated) (OR = 11.54; 95% CI: 1.18 - 112.58; P = 0.0426). This was however not significant with multivariate analysis (OR = 9.00; 95% CI: 0.89 - 90.91) (Table 5).

**Table 5.** Association between variables and knowledge of healthcare workers towards HIV palliative care in public secondary and tertiary hospitals, Abuja, Nigeria-May, 2017

| Characteristics | Good Knowledge<br>N (%) | Poor Knowledge<br>N (%) | OR (95% CI) | aOR (95% CI) |
| --- | --- | --- | --- | --- |
| **Age (years):** |  |  |  |  |
| • ≤ 40 | 25 (11.0) | 203 (89.0) | 1.22 (0.58–2.57) |  |
| • > 40 | 11 (9.2) | 109 (90.8) |  |  |
| **Sex:** |  |  |  |  |
| • Male | 15 (10.2) | 132 (89.8) | 0.97 (0.48–1.96) |  |
| • Female | 21 (10.5) | 180 (89.5) |  |  |
| **Marital status:** |  |  |  |  |
| • Married | 31 (11.3) | 243 (88.7) | 1.76 (0.66–4.70) |  |
| • Unmarried | 5 (6.8) | 69 (93.2) |  |  |
| **Profession:** |  |  |  |  |
| • Doctor | 17 (7.7) | 205 (92.3) | 2.13 (1.07–4.35) * | 2.7 (1.28 – 5.56) |
| • Nurse | 19 (15.1) | 107 (84.9) |  |  |
| **Highest academic Qualification:** |  |  |  |  |
| • Bachelor degree & above | 28 (12.1) | 203 (87.9) | 1.88 (0.83 – 4.26) |  |
| • Diploma | 8 (6.8) | 109 (93.2) |  |  |
| **Work experience (years):** |  |  |  |  |
| • ≤ 10 | 24 (10.4) | 206 (89.6) | 0.99 (0.48 – 2.06) |  |
| • > 10 | 12 (10.5) | 102 (89.5) |  |  |
| **Facility type:** |  |  |  |  |
| • Secondary | 28 (15.6) | 152 (84.4) | 3.67(1.63– 8.33) * | 5.56 (2.13 – 14.29) |
| • Tertiary | 8 (4.8) | 160 (95.2) |  |  |
| **Received undergraduate training on palliative care:** |  |  |  |  |
| • Yes | 13 (7.2) | 168 (92.8) | 0.48 (0.24 – 0.99) * | 0.73 (0.34 – 1.58) |
| • No | 23 (13.8) | 144 (86.2) |  |  |
| **Received in-service training on Palliative care** |  |  |  |  |
| • Yes | 5 (4.1) | 116 (95.9) | 0.27 (0.10- 0.72) * | 0.32 (0.11- 0.90) |
| • No | 31 (13.7) | 196 (86.3) |  |  |

\*Statistically significant at p-value ≤0.05

*Statistically significant at p-value ≤0.05

Professional qualification and highest academic qualification were positively associated with good practice while secondary health facility, undergraduate training and in-service training on palliative care were negatively associated with health care worker’s practice on HIV palliative care (Table 6)

**Table 6.** Associations between variables and practice of healthcare workers towards HIV palliative care in public secondary and tertiary hospitals in FCT, Abuja, Nigeria-May, 2017

| Characteristics | Good practice<br>N (%) | Poor practice<br>N (%) | OR (95% CI) | aOR (95% CI) |
| --- | --- | --- | --- | --- |
| **Age (years):** |  |  |  |  |
| • ≤ 40 | 66 (29.0) | 162 (71.0) | 1.63 (0.96 – 2.77) |  |
| • > 40 | 24 (20.0) | 96 (80.0) |  |  |
| **Sex:** |  |  |  |  |
| • Male | 36 (24.5) | 111 (75.5) | 0.88 (0.54 – 1.44) |  |
| • Female | 54 (26.9) | 147 (73.1) |  |  |
| **Marital status:** |  |  |  |  |
| • Married | 71 (25.9) | 203 (74.1) | 1.01 (0.56 – 1.82) |  |
| • Unmarried | 19 (25.7) | 55 (74.3) |  |  |
| **Profession:** |  |  |  |  |
| • Doctor | 50 (39.7) | 76 (60.3) | 2.99(1.83 – 4.91)* | 2.04 (1.03-4.02) |
| • Nurse | 40 (18.0) | 182 (82.0) |  |  |
| **Highest academic Qualification:** |  |  |  |  |
| • Bachelor degree & above | 74 (32.0) | 157 (68.0) | 2.98(1.64 – 5.40)* | 2.30 (1.05-5.08) |
| • Diploma | 16 (13.7) | 101 (86.3) |  |  |
| **Work experience (years):** |  |  |  |  |
| • ≤ 10 | 66 (28.7) | 164 (71.3) | 1.51 (0.89 – 2.57) |  |
| • > 10 | 24 (21.0) | 90 (79.0) |  |  |
| **Facility type:** |  |  |  |  |
| • Secondary | 20 (11.1) | 160 (88.9) | 0.18(0.10 – 0.32)* | 0.15 (0.08-0.28) |
| • Tertiary | 70 (41.7) | 98 (58.3) |  |  |
| **Received undergraduate training on palliative care:** |  |  |  |  |
| • Yes | 22 (12.1) | 159 (87.9) | 0.20 (0.12-0.35)* | 0.23 (0.12-0.44) |
| • No | 68 (40.7) | 99 (59.3) |  |  |
| **Received in-service training on Palliative care** |  |  |  |  |
| • Yes | 15 (12.4) | 106 (87.6) | 0.29 (0.16-0.53)* | 0.53 (0.25-1.11) |
| • No | 75 (33.0) | 152 (67.0) |  |  |

\*Statistically significant at p-value ≤0.05

*Statistically significant at p-value ≤0.05

## Discussion

Over half of the participants in this study were female, majority were nurses and most of them possess a minimum of bachelor degree. About half of respondents claimed they were trained in palliative care during undergraduate study period while very few had received in-service training on palliative care.

Majority of the health workers had poor knowledge of palliative care and this will adversely impact effective service delivery. This finding is consistent with those from the works of Kassa[13], Gedamu[15] and Morsy[16] but differs in some other studies where level of knowledge of participants were high[17, 18]. A possible reason for this poor palliative care knowledge could be because majority of the study participants had not had training on palliative care either as in-service or during undergraduate study[19’ 20’ 21].

It is interesting to know that participants who did not receive undergraduate training on palliative care and those without in-service training demonstrated good knowledge of palliative care better than those who had been trained respectively. This differs from most other studies where training was found to have a positive significant association with good knowledge[13]’[15] ’[18]. The variation in our findings could be due to the fact that the content and quality of the training were poor or inadequate as reported by Gedamu[15]. It’s also possible that the period between the training and the time of this assessment could result in poor recall. The implication of this is that poor, inadequate or infrequent training on palliative care could hinder effective service delivery.

Participants in secondary facilities demonstrated good knowledge of palliative care better than their contemporaries in tertiary health facilities. It is expected that good knowledge of palliative care should be higher in tertiary facility being a teaching hospital and a referral centre. This finding could have been due to the knowledge possessed by the individuals rather than the influence of the facilities. Operation based on restriction to professional speciality in tertiary health facilities could also be another possible reason for the low knowledge in the tertiary facilities unlike in the secondary where each staff are involved in almost every are of patients care. Demand from workload among tertiary health facility staff could also result into reduced their attention on subject matters.

Majority of the participants also had misconceptions about palliative. The most common ones were that palliative care (PC) “PC is disease-oriented and not person oriented”, “PC is concerned with prolongation of life”, PC is just terminal care and that “the use of placebos is appropriate in the treatment of some types of pain”. This could be a demonstration of the poor quality of the training received by participants on palliative care or could be a reflection of how people who need palliative care are being managed. This finding is similar to works done by some other researchers [15]’[16]’[16]. Poor misconception about palliative care will negatively influence how healthcare deliver to the PLWHA in need of it. Majority of participants believed that people living with HIV/AIDS actually requires palliative care. This shows there is an understanding of the expansion of the scope of palliative from caring for cancer patients to involving HIV infection and other noncommunicable diseases[22].

Regarding the attitude of healthcare workers towards palliative care, almost all the health care workers had favourable attitude despite a lack of formal and in-service training of most of the staff. Though the proportion of attitude is higher than most other studies, the finding is still in keeping with others studies conducted in India,[12]’[23] and Ethiopia [17]’[15] & [13]. However, other studies showed that less than half of the health care workers have unfavourable attitude towards palliative care[18]. Our finding might possibly be influenced by the FATCOD scale score used for determination of good and poor attitude. It could also be as a result of the natural passion and the training that comes with the health profession.

Participants who were married were found to possess a more favourable attitude towards palliative care for PLWHA though this was not significant during multivariate analysis. This is also similar to the findings in the works done in Ethiopia[13]. Favourable attitude towards palliative care will lead to improved service provision for PLWHA which will also improve their disease outcome. However, training, length of work experience, academic qualification and professional discipline which have been known to influence attitude were not significant in this study. All this could have been influenced by poor knowledge of palliative care.

About half of the respondents were in agreement that palliative care should be given to “only dying PLWHA”. This is a misconception as it is expected that palliative care should be given to every person living with HIV/AIDS. The misconception could be due to the believe that palliative care is the same as end-of-life care. Agreement to this misconception is higher than the findings from other studies where only 15.3%[13], 21.4%[15] and 33.4%[15] are in agreement with this attitude [13],[15].

Over half of the participants said they were uncomfortable with talking about death with a dying PLWHA. This is similar to the study done by Gedamu [15] but higher than other studies done in Egypt [24] and South Africa [25]. This might be associated with our culture, tradition or religious factors that make brings about reluctance in discussing the issue of death with patients. About one-third of health care workers would not want to be assigned to a dying PLWHA. This is lower than the findings from previous studies[13]’[24, 25]. Feeling of incompetence in providing the necessary care for the patients may be responsible from our findings.

About three-quarters of the respondents believe that it is best to change the subject to something cheerful when an HIV/AIDS patient asks “am I dying?” Agreement on this attitude is higher than from other studies.[13]’[15]. This again may not be unrelated to religious or cultural issues influencing breaking of bad news to patients. It is advisable to tell dying patients the truth about their condition so that they can make necessary preparation with regards to family, legal and religious issues [26]. as hiding the truth from patients has been considered unethical [27].

Regarding the practice of palliative care for PLWHA, about three-quarter of the participants had poor practice towards palliative care. This level of poor palliative care practice indicates that the quality of palliative care services offered to PLWHA could be very poor. Our finding is similar with study conducted by Kassa where most participants were reported to have poor practice with regards to palliative care[13]. This finding could be related to the poor knowledge of participants about palliative care especially as knowledge correlates positively with practice.

Determinants of good practice of palliative care were being a doctor, possessing a minimum of bachelor degree and working in tertiary health facilities. However, participants whose undergraduate curriculum did not contain palliative care or who have not had any in-service training on palliative care were found to demonstrate good practice when compare with others. These differ from the earlier studies where those who had training on palliative care both at undergraduate and formal inservice training demonstrate better practice of palliative care than those who had not been trained[18]. This finding might be due because the contents of training were inadequate or recall was poor due to the interval between their training and when our study what was taught.

Most participants would inform terminally ill patients and PLWHA about their diagnosis and this is much higher compare to previous studies in Ethiopia and Lebanon[13][28][18]. This might be due to the fact that both doctors and nurses have the responsibility of discussing patient’s diagnosis with them unlike in Ethiopia where it is mainly the responsibility of the doctors[13]. Majority considers medical and about half of the respondents consider spiritual conditions when dealing with terminally ill patients or PLWHA while about two-third connect with spiritual counsellor when handling spiritual matters. This could be due to the Nigerian’s value for religious belief. This is similar to findings from Lebanon and Ethiopia[13][28].

Morphine and Pentazocine were the most commonly used drugs by respondents for the treatment of severe pain unlike the study in Ethiopia where Paracetamol or ibuprofen were the most commonly used drugs for such[13]>[18]. Our finding could be based on healthcare workers’ effort to relief patients’ pain by all means or lack of understanding of WHO analgesic ladders use. However, if appropriately used to relief pain, patients’ quality of life will improve[13]. More so, use of these opioids could also be due to increased access to drugs as there were no restriction to morphine for cancer patients[29] unlike the restriction placed on opioids in Ethiopia[13].

The study was conducted among doctors and nurses who play important roles in the delivery of palliative care to PLWHA. However, new or broader input from the respondents were precluded because a closed-ended question were used in the study instruments.

## Conclusion

Majority of the respondents had favourable attitude towards palliative care despite a poor knowledge and practice with regards to palliative care for people living with HIV/AIDS. Poor knowledge and practice of palliative care among healthcare workers will negatively impact the quality of service delivery of palliative care to PLWHA. We recommended that quality in-service training and continuous education on palliative care should be regularly given to improve doctors and nurses’ knowledge and practice.

## Acknowledgements

We acknowledge the data team of the African Field Epidemiology Network (AFENET). We would also like to express our profound gratitude to the study participants and our data collectors without whom our study could be terminated halfway into our task.

